# Diversity reduces extinction risk at global scales

**DOI:** 10.1101/2020.09.17.296285

**Authors:** Brian C. Weeks, Shahid Naeem, Jesse R. Lasky, Joseph A. Tobias

**Author notes:** **Statement of authorship:** BW conceived the study and performed the analyses; JT and SN helped develop the conceptual framework; JT provided trait data; JR contributed to the analyses; all authors contributed to writing and revising the manuscript. **Data accessibility statement:** Trait data used in this study are published openly in the same volume, and the phylogenetic and distributional data are publicly available.

## Abstract

Increases in biodiversity often lead to greater, and less variable, levels of ecosystem functioning. However, whether species are therefore less likely to go extinct in more diverse systems is unclear. We use comprehensive estimates of avian taxonomic, phylogenetic and functional diversity to characterize the global relationship between multiple dimensions of diversity and extinction risk in birds. We find that more diverse assemblages have lower mean IUCN threat status despite being composed of species with attributes that make them more vulnerable to extinction, such as large body size or small range size. Our analyses also reveal that this reduction of current threat status associated with greater diversity far outweighs the increased risk associated with the accumulation of extinction-prone species in more diverse assemblages. These results suggest that species conservation targets can best be achieved by maintaining high levels of overall biodiversity in natural ecosystems.

## Introduction

Numerous experimental and observational studies have shown that biodiversity is positively associated with an array of ecosystem functions (Cardinale *et al*. 2002, 2006; Emmett Duffy *et al*. 2017). Increasingly, research on biodiversity–ecosystem function (BEF) relationships is revealing that diversity-driven increases in function can boost rates at which nutrients, energy and organic matter flow through an ecosystem (Cardinale *et al*. 2012), as well as increasing its overall multifunctionality (Soliveres *et al*. 2016), stability (Tilman *et al*. 2014) and resilience (Oliver *et al*. 2015). In addition, increased diversity is associated with reduced rates of species invasion (Naeem *et al*. 2000; Levine *et al*. 2004; Fargione & Tilman 2005; Byun *et al*. 2013) and lower rates of disease transmission (Becker *et al*. 2014). These benefits are generally conceptualized at the scale of whole ecosystems, yet it is also possible that they influence the fate of individual lineages by reducing extinction risk (Weeks *et al*. 2016b). However, the relationship between the diversity of an assemblage and the risk of extinction for its constituent lineages is rarely investigated and remains poorly understood.

A key hindrance to progress is that this question is unlikely to be resolved when biodiversity is measured simply in terms of species richness (i.e. taxonomic diversity). Extinction risk may be more closely associated with other aspects of ecosystems, including functional and phylogenetic components of biodiversity (Naeem *et al*. 2016). For example, functional traits often perform better than species richness in predicting ecosystem function and stability (Tilman *et al*. 1997; Hooper *et al*. 2005), suggesting that extinction risk may be more sensitive to variation in functional diversity. Accounting for the multidimensionality of diversity is also important because different facets of biodiversity can have contrasting responses to environmental change (Chapman *et al*. 2018) and vary in their predicted relationships with ecosystem function, as well as the mechanisms underpinning those relationships (Flynn *et al*. 2011; Soliveres *et al*. 2016). As yet, it has not been possible to account for such multidimensionality in studies of extinction risk because the necessary combination of species-level information on geographical distributions, phylogenetic relationships and detailed functional traits have not generally been available at sufficiently large spatial and taxonomic scales (Naeem *et al*. 2016).

Capitalizing on the availability of comprehensive phylogenetic (Jetz *et al*. 2012) and distributional data for birds (BirdLife International 2015), we develop a multidimensional metric of avian diversity to explore its association with extinction risk at a global scale. Birds offer an ideal system for this approach because they are distributed worldwide with high quality species-level information on co-occurrence, threat status and—increasingly—functional traits (Tobias *et al*. 2020). Using a newly compiled data set of morphological trait measurements from >40,000 individual birds of >10,000 species, representing >99% of bird species diversity (Tobias *et al*., this issue), we calculate functional richness (Villéger *et al*. 2008) for avian assemblages based on body mass, beak shape, leg length and tail length. At global scales, these traits provide a powerful index of avian dietary niche and foraging behaviour (Pigot *et al*. 2020). Our estimation of functional richness therefore focuses on ‘effect traits’ (i.e. traits that determine the contribution of an individual to ecosystem functioning; Winemiller *et al*. 2015).

Since eco-morphological and life history traits are also linked to conservation status in birds (Tobias & Pigot 2019), we use them to develop a metric of extinction risk. We assume that increases in body mass and ecological specialization, as well as decreases in geographical range size and dispersal ability, are associated with the increased likelihood that a lineage will go extinct per unit time, as reported in numerous studies (e.g. Bennett & Owens 1997; Sekercioglu *et al*. 2004; Reinhardt *et al*. 2005; Lee & Jetz 2011; Jetz & Freckleton 2015). We quantify dispersal ability using wing morphology (Claramunt *et al*. 2012; Sheard *et al*. 2020), and we estimate specialization based on the trophic diversity of each species’ diet (Wilman *et al*. 2014; Pigot *et al*. 2020). Because these attributes predict the probability that a species will go extinct, we use our trait-based metric of extinction risk to calculate the collective vulnerability of species in assemblages, or ‘assemblage vulnerability’ (Weeks *et al*. 2016b). In other words, assemblages composed of species with small range sizes, low dispersal abilities, large body sizes and high levels of ecological specialization have greater overall vulnerability. Since our calculation of assemblage vulnerability is partly based on the presence of species not currently considered threatened with extinction, but likely to become threatened in the future, it provides a measure of latent extinction risk (i.e., the difference between a species’ contemporary extinction risk, and the expected level of risk, given its biology; Cardillo *et al*. 2006).

Although they can theoretically capture collective or latent extinction risk, trait-based metrics provide a relatively crude estimate of contemporary extinction risk (Tobias & Pigot 2019). Thus, we also characterize the mean threat status of assemblages using IUCN Red List status (BirdLife International 2015), an indicator of current conservation priorities widely used in global-scale analyses (Isaac et al. 2007). Previous studies have shown that IUCN Red List status and trait-based predictors of extinction risk are correlated in birds (Tobias & Pigot 2019), but it is less clear how they are linked to biodiversity. Although the standard prediction based on BEF literature is that biodiversity enhances ecosystem functioning, thereby reducing extinction risk, other factors may complicate the outcome. In particular, if occurrence within diverse assemblages reduces rates of extinction for individual lineages, this may—paradoxically— increase assemblage vulnerability through the survival and accumulation of extinction-prone species (Weeks *et al*. 2016b; Fig. 1). These contrasting possibilities set up a potential trade-off whereby increased diversity may have both positive and negative implications from the perspective of biological conservation.

**Figure 1.**
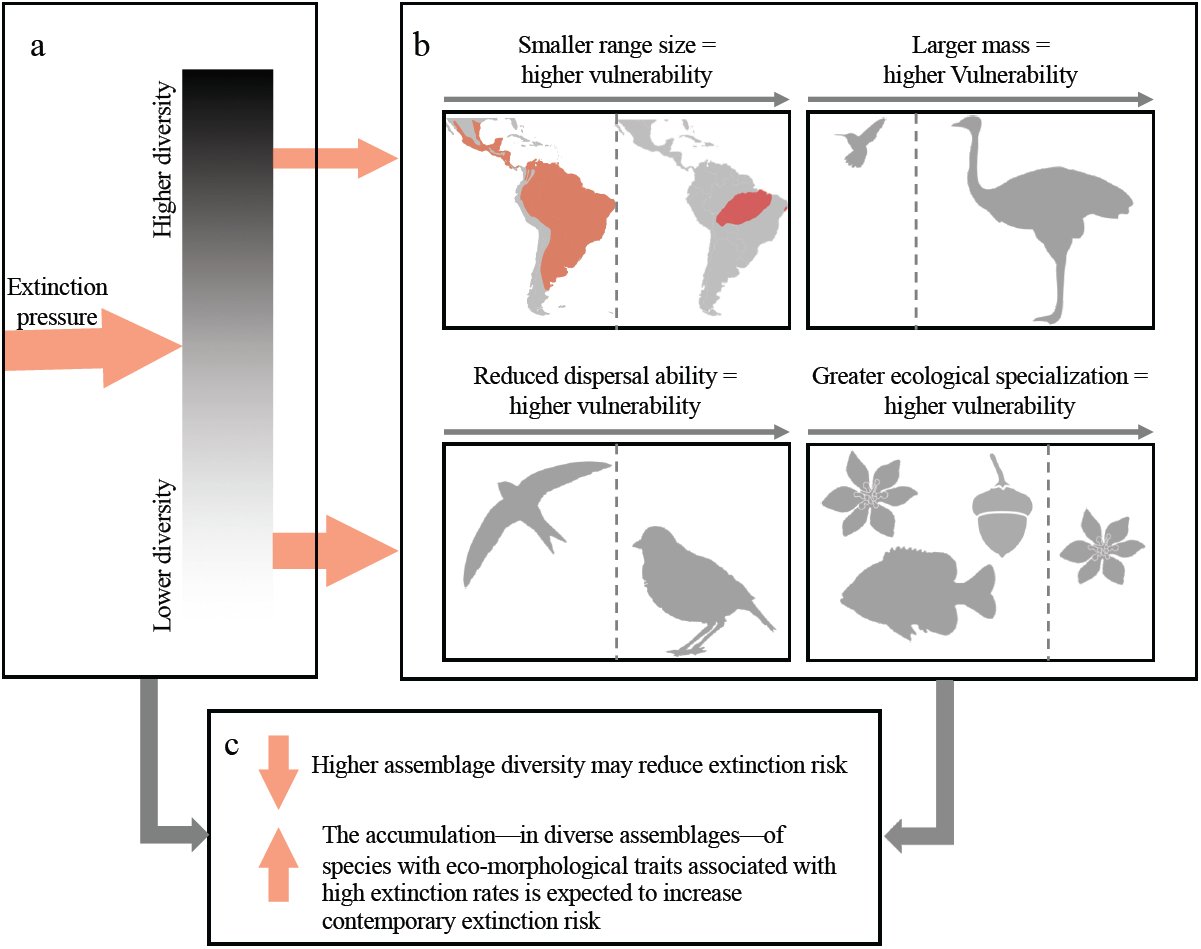
Conceptual illustration of the relationship between diversity and extinction risk. (a) Species in more diverse assemblages are hypothesized to have reduced exposure to extinction pressure as a result of biodiversity effects on ecosystem functioning and stability. (b) The phenotypic and biogeographic attributes of individual species in an assemblage determine the impacts of the extinction pressures to which they are exposed (i.e. their vulnerability). (c) Together, the diversity and attributes of constituent species within an assemblage determine the contemporary extinction risk of assemblages. Thus, the relationship between diversity and extinction risk may depend on a trade-off between two inter-dependent processes: 1) the reduction of extinction risk associated with higher assemblage diversity (a→c), and 2) the consequent accumulation of vulnerable species in more diverse assemblages (a→b→c).

Comparing across taxonomic, phylogenetic and functional diversity metrics, we examine the effects of different components of bird diversity on assemblage vulnerability and IUCN threat status (Fig. 2). In addition, to shed light on underlying processes, we use structural equation modeling to quantify the strength of the relationships between bird diversity, assemblage vulnerability and extinction risk. The findings allow us to disentangle the positive and negative effects of biodiversity on contemporary and latent extinction risk, with implications for the prioritization of conservation interventions.

**Figure 2.**
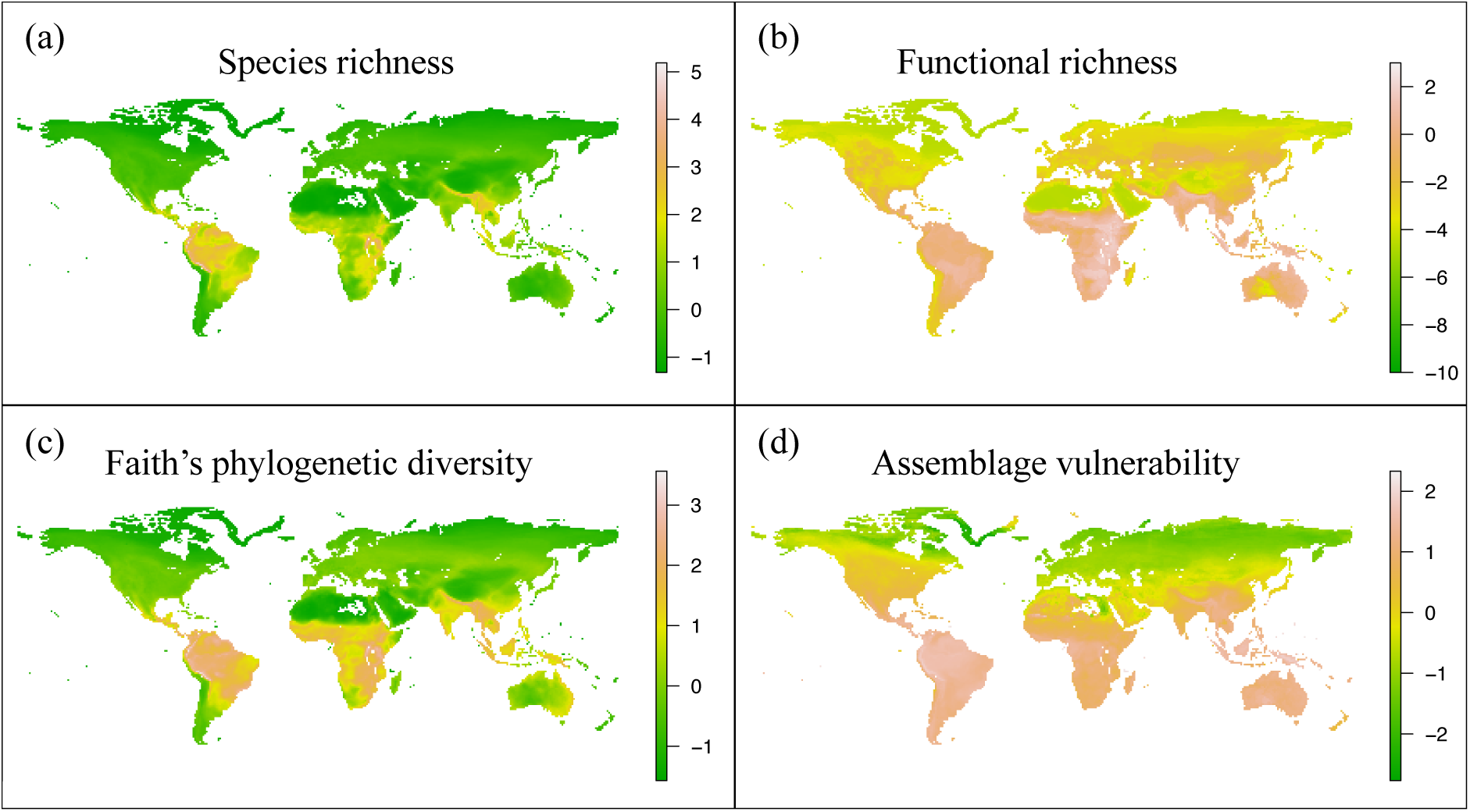
Relationship between bird diversity and assemblage vulnerability mapped at global scales. Patterns shown are based on estimates of (a) taxonomic, (b) functional, and (c) phylogenetic diversity calculated from all species mapped as occurring in 1 degree grid cells worldwide. Functional diversity is estimated from morphological traits for over 10,000 bird species. We use biogeographical, ecological and morphological attributes of all species to generate an index of assemblage vulnerability (a metric of mean vulnerability to extinction for species in an assemblage). Maps show each variable standardized to a mean of 0 and standard deviation of 1; the logarithm of standardized functional richness estimates that were transformed to be positive are mapped.

## Methods

### Presence-Absence Matrix

We developed a 1° latitude by 1° longitude resolution occurrence map for terrestrial bird communities, excluding all non-terrestrial cells (those that were > 50% ocean or > 70% inland water), and all cells below 60° S. We then determined species composition of these cells using species range maps at a 102 km resolution, obtained from BirdLife International, projected onto a 1° latitude by 1° longitude grid. Species ranges were trimmed to exclude areas where presence was classified as uncertain or extinct. We also omitted areas where species origin was classified as vagrant or uncertain, and where seasonality was classified as passage (i.e. only occurring on migration) or uncertain. Species were considered to be present in a grid cell assemblage if their range covered at least 10% of a cell (unless a species range was sufficiently small that it did not cover 10% of any cell, in which case that species was considered to be present in any cell that overlapped with their range). Any cells with fewer than 7 species were removed, so that each cell had enough taxa to calculate functional richness using 6 traits (Villéger *et al*. 2008). While species occurring in the same 1° grid cell do not necessarily interact as a community, the total avian assemblage of each cell serves as an estimate of the complete range of traits and trophic interactions that could potentially contribute to ecological functions with relevance to extinction risk. At global scales, quantification and validation of interspecific interactions is not feasible, so co-occurrence within 1° cells is routinely used as a proxy for coexistence (e.g. Pigot *et al*. 2016) or to link biodiversity and ecosystem function (e.g. Duchenne *et al*. 2020).

### Community Diversity Metrics

For each assemblage occupying each grid cell, we calculated species richness, two measures of phylogenetic diversity, and one metric of functional diversity (Fig. 2). Functional diversity was characterized using six ecologically-important functional effect traits (total beak length, beak tip to the anterior edge of the nares, beak width, beak depth, tail length, and tarsus length) measured on museum specimens (Pigot *et al*. 2020). For each assemblage, we used these traits to calculate functional richness—the volume of the convex hull that bounds the functional trait space (Villéger *et al*. 2008)—using the ‘dbfd’ function in the FD R package (Laliberté & Legendre 2010; Laliberté *et al*. 2015; R Core Team 2018). All traits were standardized to a mean of zero and standard deviation of one prior to analysis. In the ‘picante’ package in R (Kembel *et al*. 2010), we used the ‘pd’ and ‘cophynetic’ functions, respectively, to calculate the phylogenetic diversity of each assemblage as (1) the sum of the branch lengths connecting all species in the community—i.e. Faith’s phylogenetic diversity index (Faith 1992)—and (2) the mean pairwise phylogenetic distance (Webb *et al*. 2002) between all species in the community.

The phylogenetic relationships among species were estimated using 1,000 phylogenies taken from the posterior distribution of the Jetz et al. (2012) global phylogeny of birds, with the (Hackett *et al*. 2008) phylogeny used as a backbone, and including species that were placed by Jetz et al. (2012) using taxonomy. From these phylogenies, we calculated a maximum credibility clade tree using DendroPy (Sukumaran & Holder 2010) as described in Rubolini *et al*. (2015).

### Assemblage Vulnerability

To calculate the accumulation of species with traits and distributions that make them pre-disposed to extinction, we quantified assemblage vulnerability for each assemblage in the world, based on a modification of the approach taken by Weeks *et al*. (2016b). All variables were standardized to a mean of zero and standard deviation of one prior to calculation of vulnerability for both species and assemblages. For each species in an assemblage, we calculated a species-specific trait-based vulnerability score (V_t_, eqn 1) based on dispersal ability (measured as hand-wing index (Claramunt *et al*. 2012) with data from (Sheard *et al*. 2020)), mass from Tobias & Pigot (2019), and the degree of specialization (based on the trophic diversity of their diets (Wilman *et al*. 2014; Pigot *et al*. 2020):

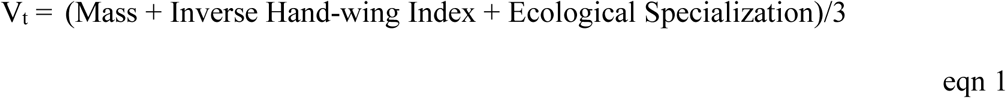

We then considered the mean of V_t_ and the inverse of range size (which we considered to be the number of cells in which a species was present) to be each species’ vulnerability (V_s_):

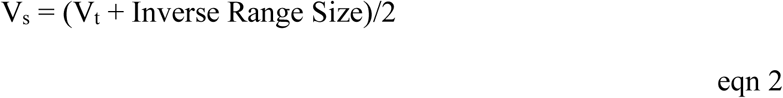

We then calculated the assemblage vulnerability for each assemblage, ‘i’, as the unweighted mean of the vulnerability scores (V_s_) for all (n) species in an assemblage:

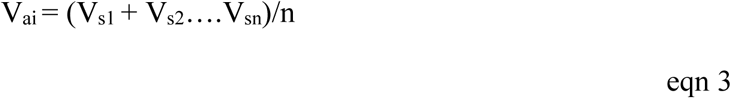

Thus, larger species with low dispersal abilities, greater ecological specialization, and smaller range sizes would have higher species vulnerability (V_s_) scores, and assemblages that are composed of species that tended to have higher V_s_ scores would have higher assemblage vulnerability (V_a_) scores.

### Contemporary Threat Status

To characterize the contemporary threat status of each assemblage, we calculated the mean IUCN threat status (BirdLife International 2015) of all species in the assemblage. We converted threat status categories to numeric variables, a commonly used approach for characterizing extinction risk (Isaac *et al*. 2007), with Least Concern = 1 and Critically Endangered = 5. For each assemblage, we then calculated the mean IUCN threat status of its constituent species, and standardized the assemblage-level variable to have a mean of 0 and a standard deviation of 1, to improve model fitting.

### Structural Equation Modeling

We characterized diversity as a latent variable reflected in the observed (i.e. exogenous, as opposed to latent) covariates: species richness, functional richness, phylogenetic diversity, and mean pairwise phylogenetic distance measures of the species in an assemblage (Fig. 3). This approach is based on the conceptual framework of Naeem *et al*. (2016) in which diversity is treated as a multidimensional construct, with each exogenous predictor measured as described in the *Community Diversity Metrics* section, above. The loading of species richness on diversity was set to 1 in order to define the scale of the latent diversity variable (Rosseel 2012). We then modeled assemblage vulnerability as a function of diversity, and contemporary threat status as a function of diversity and assemblage vulnerability (Fig. 3). Each path coefficient linking two variables was considered to be the direct effect of the predictor variable on the response. The indirect effect of diversity on contemporary threat status (via the effect of diversity on assemblage vulnerability) was calculated as the product of the path coefficient linking diversity and assemblage vulnerability and the path coefficient linking assemblage vulnerability and contemporary threat status. All reported coefficients are standardized.

**Figure 3.**
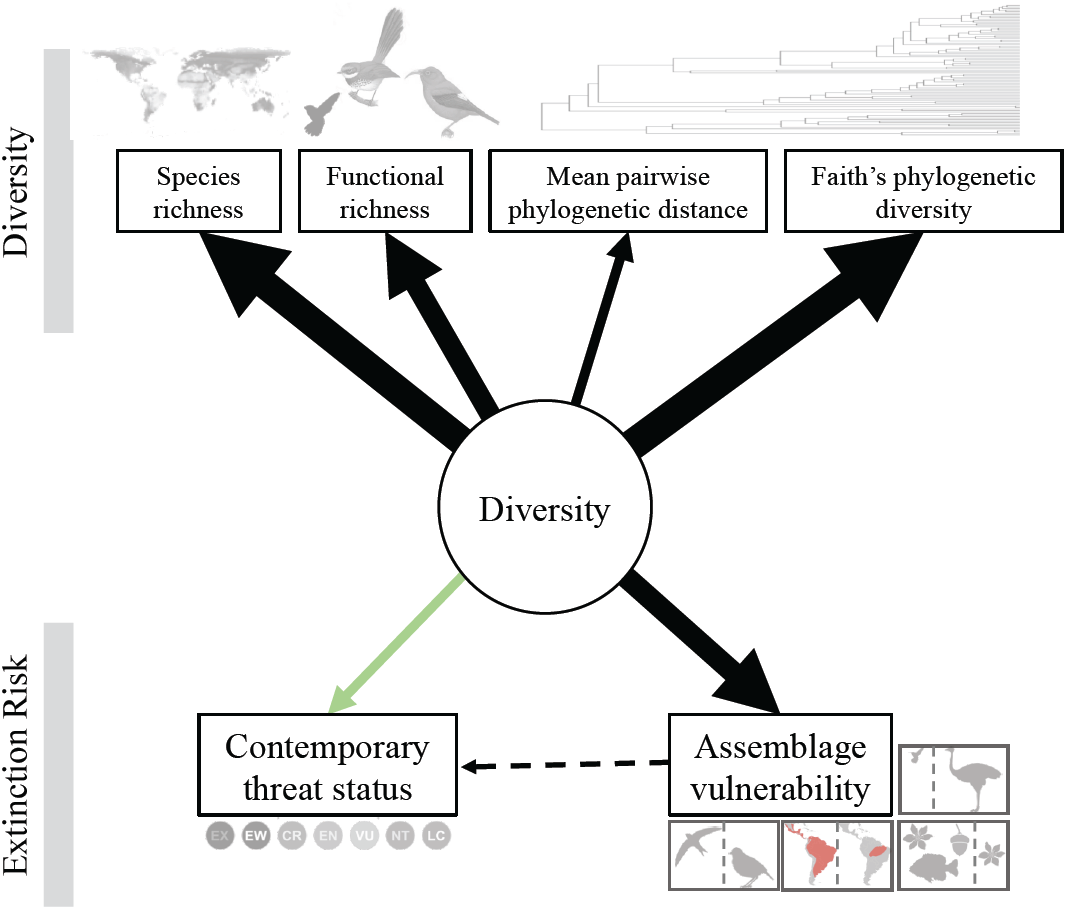
Diversity influences extinction risk in birds. We use structural equation modeling to explore links between bird diversity, assemblage vulnerability, and contemporary threat status. Black arrows indicate positive relationships; the green arrow indicates a negative relationship; all relationships are significant. The width of the arrows is scaled to standardized effect size, solid lines represent significant relationships and dashed lines represent relationships that are not significant. We model bird assemblage diversity as a latent variable, measured using alternative univariate dimensions (taxonomic, functional and phylogenetic diversity). The arrows connecting diversity to univariate dimensions represent the loadings of diversity on each dimension. Arrows connecting diversity, assemblage vulnerability, and contemporary threat status represent regression parameters connecting the predictor to response variables. Diversity has a strong positive relationship with assemblage vulnerability, suggesting that species predisposed to extinction have accumulated in diverse assemblages. Via this pathway, diversity has a positive effect on contemporary threat status although, importantly, this is an order of magnitude weaker than the direct negative effect of diversity, and this relationship is not significant (i.e. diversity reduces contemporary threat status overall).

To account for the potential impacts of spatial autocorrelation, we fit the structural equation model using a spatially explicit structural equation modeling approach (Rosseel 2012; Lamb *et al*. 2014). Because the size of the dataset (15,244 assemblages) precluded the calculation of a comprehensive spatial distance matrix, we randomly subsampled 1,000 assemblages from the dataset without replacement and fit the model to the subset of the data. To account for spatial autocorrelation, we fit the SEM using 10 lag distances, with an upper limit of 50% of the total distance between subsampled points, using the ‘runModels’ function from the SESEM package in R (Lamb *et al*. 2014; R Core Team 2018). This returns 10 fitted SEMs, which we compared using the comparative fit index (CFI). We then extracted the model statistics, parameters, and parameter significance estimates from the SEM based on the lag distance that produced an SEM with the highest CFI value.

We repeated this model fitting on random subsets of the data 1,000 times and calculated the mean parameter estimate for each path across all random subsets. We considered the 2.5%-97.5% quantiles to be the confidence interval (CI) of each parameter, and considered parameters to be significant if the CI did not overlap with zero. We assessed overall model fit using the comparative fit index (CFI) and root mean square error of approximation (RMSEA).

## Results

We characterized functional, phylogenetic, and taxonomic diversity for 15,224 avian assemblages worldwide (Fig. 2). In our model, our latent diversity variable had a variance of 1, and positive loadings on all exogenous predictors of diversity: species richness (*β* = 0.96), functional richness (*β* = 0.69), Faith’s phylogenetic diversity (*β* = 1.03), mean pairwise phylogenetic distance (*β* = 0.32), and all relationships between these exogenous variables and diversity were significant (Fig. 3).

We report average model fit statistics from across 1,000 model fits from random subsets of 1,000 assemblages from our dataset. While metrics of the absolute goodness of fit were relatively low (RMSEA = 0.15, *P* < 0.01; χ^2^ = 7,524, df = 8, *P* < 0.01), this is not particularly surprising given the scope of the dataset, the simplicity of the model, and the tendency for ecological data to be noisy. More importantly, the model had a better fit to the data than a null model: CFI = 0.97, CI = 0.96 – 0.98, with anything over CFI = 0.90 implying a good fit to the data (Kline 2010).

We found that diversity was significantly positively associated with assemblage vulnerability (*β* = 0.72, CI = 0.68 - 0.76), and also explained a substantial fraction of the variance in assemblage vulnerability (assemblage vulnerability *R*2 = 0.52), indicating that more diverse assemblages were on average more vulnerable (i.e. had higher V_s_; eqn 2), associated with latent extinction risk. Conversely, diversity was significantly negatively associated with contemporary extinction risk, indicating more diverse assemblages had lower mean IUCN threat level (*β* = - 0.31, CI = -0.44 – -0.17). Assemblage vulnerability was positively associated with contemporary threat status, but the effect size was small and the relationship was not significant (*β* = 0.03, CI = -0.17 – 0.23). As a result, while diversity had a significant negative direct effect on contemporary extinction risk (*β* = -0.31), it had a contrasting positive indirect effect on contemporary extinction risk (*β =* 0.02; Fig. 3), but this relationship was not significant. Thus, the indirect negative effect of diversity on contemporary threat, driven by the accumulation of more vulnerable species, may have the potential to partly limit the benefit of diversity in reducing contemporary threats, but this contrasting effect is relatively small and non-significant. The model explained roughly 9% of the variance in contemporary threat status (*R*2 = 0.09, CI = 0.06 – 0.13).

## Discussion

By compiling multiple dimensions of diversity data for the global avifauna, we have shown that species occurring in assemblages with higher levels of diversity have reduced contemporary extinction risk. It may seem intuitive that reduced extinction risk has resulted in increased diversity, particularly over deeper timescales at which declining extinction rates towards the equator have allowed species richness to build up in tropical biota, driving latitudinal diversity gradients (Mittelbach *et al*. 2007). However, our analyses focus on contemporary and latent extinction risk, a temporal scale less relevant to the effect of diversification or glaciation, and more relevant to the near-term trends determining IUCN Red List status and vulnerability.

Recent anthropogenic threats have driven relatively few bird lineages to extinction, but have caused a significant proportion of global avian diversity to be classified as threatened (BirdLife International 2015). At this temporal scale, our results are more likely to be explained by characteristics of diverse ecosystems, with increases across multiple facets of diversity reflecting a higher level and stability of ecosystem functioning. This may take the form of more complete networks of species interactions and associated processes, or the buffering effect of biodiversity against risks such as invasion or disease (Naeem *et al*. 2000; Levine *et al*. 2004; Fargione & Tilman 2005; Byun *et al*. 2013; Becker *et al*. 2014).

While the direct reduction in contemporary extinction risk associated with diversity suggests that species in more diverse assemblages are at lower risk of extinction, this relationship is complicated by the dynamic history of community assembly (Weeks *et al*. 2016a). For example, reduced extinction pressure may result in the long-term survival of species otherwise prone to extinction, which therefore tend to accumulate in diverse assemblages over time (Weeks *et al*. 2016b). There is some evidence that this occurs in plants: climatic stability is thought to have reduced extinction risk for rare species, allowing them to persist in climatically stable regions, with the result that climate change and anthropogenic drivers of extinction are now disproportionately impacting rare species in more diverse regions (Enquist *et al*. 2019).

In accordance with the idea that diversity can both decrease short-term and increase long-term vulnerability, we find that the reduction in contemporary extinction risk (*β* = -0.31) is coupled with increased latent extinction risk as measured by assemblage vulnerability (*β* = 0.72). This suggests that more diverse communities are composed of many species that are not currently threatened but with attributes associated with higher risk of extinction: poor dispersal ability, large body size, greater ecological specialization, and smaller range sizes. One possible interpretation of this pattern is that attributes associated with increased vulnerability may promote diversification (e.g., reduced dispersal ability can lead to increased diversification rates; Weeks & Claramunt 2014). However, the association between our indices of vulnerability and diversification at global scales is weak and mixed (Owens *et al*. 1999; Tobias *et al*. 2020), suggesting that their role as drivers of diversification is unlikely to explain our results. Overall, we interpret the elevated vulnerability of diverse assemblages as an outcome of lower rates of extinction for extinction-prone species, suggesting that the long-term consequence of lower contemporary extinction risk is an increase in latent extinction risk.

To understand the overall relationship between biodiversity and extinction risk in natural systems, it is therefore important to disentangle the contrasting effects of diversity on the current survival prospects of individual lineages (reduced short-term risk) from the accumulation of species inherently predisposed to extinction in future (increased long-term risk). When we assess the relationship between assemblage vulnerability and contemporary extinction risk, we find a slight positive association. While this result hints at an indirect mechanism by which biodiversity could ultimately be associated with increased contemporary extinction risk, the relationship between assemblage vulnerability and contemporary extinction risk is relatively weak (*β* = 0.03) and not significant. Moreover, the increase in contemporary extinction risk associated with increased assemblage vulnerability in more diverse assemblages (*β* = 0.02) is an order of magnitude weaker than the direct effect of high diversity in reducing contemporary extinction risk (*β* = -0.31). In other words, the effect of diversity in boosting latent extinction risk is negligible in comparison with its effect in reducing contemporary extinction risk.

Although the variables we use to determine assemblage vulnerability and IUCN threat status are widely considered to be indicators of extinction risk, their connection to extinction rate is complicated (Harcourt 2005). For example, some forms of rarity—beyond restricted range size (Rabinowitz 1981)—might influence IUCN threat status designations without necessarily being related to extinction rates (Harnik *et al*. 2012). While directly examining the relationship between diversity and extinction rates would be preferable, doing so is not currently feasible given our knowledge of taxa that have gone extinct is highly incomplete, particularly in relation to their ranges, traits and phylogenetic relationships. Our indices of contemporary extinction risk (IUCN threat status) and assemblage vulnerability may be poor predictors of extinction rates across timescales, which may partially explain the weak and non-significant relationship between assemblage vulnerability and contemporary threat status (Fig. 3).

The effects of biodiversity on ecosystem function can be complicated by assembly history (Fukami & Morin 2003) and temporal scale (Reich *et al*. 2012). For similar reasons, historical biogeography can alter the relationship between biodiversity and vulnerability (Weeks *et al*. 2016b). Predicting the effects of future biodiversity loss on ecosystem functioning, and thus threat status, may be further complicated by shifts in the species-specific functioning or abundance of surviving taxa (De Laender *et al*. 2016). Thus, the balance between diversity-driven reductions in contemporary extinction risk and increases in the number of species inherently sensitive to extinction may be altered according to context, with some diverse communities having higher vulnerability than others as a result of the phenotypic, biogeographic and functional attributes of their constituent species. Future studies should therefore analyze the relationship between diversity and extinction risk in different historical contexts and across a range of spatial and temporal scales.

## Conclusions

Considering spatial variation in multiple dimensions of diversity at a global scale, we find that higher diversity is associated with reduced contemporary extinction risk and increased assemblage vulnerability in birds. We attribute this general pattern to higher levels of ecosystem functioning in more biodiverse assemblages, which can theoretically moderate immediate extinction risks while also inflating the number of extinction-prone species in a community. We also show that the reduction of extinction risk associated with increased diversity is far stronger than the contrasting increase of risk associated with greater assemblage vulnerability in more diverse assemblages. We conclude that the maintenance of biodiverse communities may be a cost-effective approach to preventing extinction, reducing the longer-term need for expensive single-species conservation interventions.

## References

Becker, G.C., Rodriguez, D., Felipe Toledo, L., Longo, A.V., Lambertini, C., Correa, D.T., et al. (2014). Partitioning the net effect of host diversity on an emerging amphibian pathogen. Proc. R. Soc. B Biol. Sci., 281, 20141796.

Bennett, P.M. & Owens, I.P.F. (1997). Variation in extinction risk among birds: chance or evolutionary predisposition? Proc. R. Soc. B Biol. Sci., 264, 401–408.

BirdLife International. (2015). IUCN red list for birds. Available at: http://www.birdlife.org. Last accessed 1 January 2015.

Byun, C., de Blois, S. & Brisson, J. (2013). Plant functional group identity and diversity determine biotic resistance to invasion by an exotic grass. J. Ecol., 101, 128–139.

Cardillo, M., Mace, G.M., Gittleman, J.L. & Purvis, A. (2006). Latent extinction risk and the future battlegrounds of mammal conservation. Proc. Natl. Acad. Sci. U. S. A., 103, 4157–4161.

Cardinale, B.J., Duffy, J.E., Gonzalez, A., Hooper, D.U., Perrings, C., Venail, P., et al. (2012). Biodiversity loss and its impact on humanity. Nature, 486, 59–67.

Cardinale, B.J., Palmer, M.A. & Collins, S.L. (2002). Species diversity enhances ecosystem functioning through interspecific facilitation. Nature, 415, 426–429.

Cardinale, B.J., Srivastava, D.S., Duffy, J.E., Wright, J.P., Downing, A.L., Sankaran, M., et al. (2006). Effects of biodiversity on the functioning of trophic groups and ecosystems. Nature, 443, 989–992.

Chapman, P.M., Tobias, J.A., Edwards, D.P. & Davies, R.G. (2018). Contrasting impacts of land-use change on phylogenetic and functional diversity of tropical forest birds. J. Appl. Ecol., 55, 1604–1614.

Claramunt, S., Derryberry, E.P., Remsen Jr, J. V & Brumfield, R.T. (2012). High dispersal ability inhibits speciation in a continental radiation of passerine birds. Proc. R. Soc. B Biol. Sci., 279, 1567–1574.

Duchenne, F., Thébault, E., Michez, D., Elias, M., Drake, M., Persson, M., et al. (2020). Phenological shifts alter the seasonal structure of pollinator assemblages in Europe. Nat. Ecol. Evol., 4, 115–121.

Emmett Duffy, J., Godwin, C.M. & Cardinale, B.J. (2017). Biodiversity effects in the wild are common and as strong as key drivers of productivity. Nature, 549, 261–264.

Enquist, B.J., Feng, X., Boyle, B., Maitner, B., Newman, E.A., Jørgensen, P.M., et al. (2019). The commonness of rarity: global and future distribution of rarity across land plants. Sci. Adv., 5, eaaz0414.

Faith, D.P. (1992). Conservation evaluation and phylogenetic diversity. Biol. Conserv., 61, 1–10.

Fargione, J.E. & Tilman, D. (2005). Diversity decreases invasion via both sampling and complementarity effects. Ecol. Lett., 8, 604–611.

Flynn, D.F.B., Mirotchnick, N., Jain, M., Palmer, M.I. & Naeem, S. (2011). Functional and phylogenetic diversity as predictors of biodiversity-ecosystem-function relationships. Ecology, 92, 1573–81.

Fukami, T. & Morin, P.J. (2003). Productivity-biodiversity relationships depend on the history of community assembly. Nature, 424, 423–426.

Hackett, S.J., Kimball, R.T., Reddy, S., Bowi, R.C.K., Braun, E.L., Braun, M.J., et al. (2008). A phylogenomic study of birds reveals their evolutionary history. Science, 320, 1763–1768.

Harcourt, A.H. (2005). Problems of studying extinction risks. Science, 310, 1276.

Harnik, P.G., Simpson, C. & Payne, J.L. (2012). Long-term differences in extinction risk among the seven forms of rarity. Proc. R. Soc. B Biol. Sci., 279, 4969–4976.

Hooper, D.U., Chapin III, F.S. & Ewel, J.J. (2005). Effects of biodiversity on ecosystem functioning: a consensus of current knowledge. Ecol. Monogr., 75, 3–35.

Isaac, N.J.B., Turvey, S.T., Collen, B., Waterman, C. & Baillie, J.E.M. (2007). Mammals on the EDGE: conservation priorities based on threat and phylogeny. PLoS One, 2, e296.

Jetz, W. & Freckleton, R.P. (2015). Towards a general framework for predicting threat status of data-deficient species from phylogenetic, spatial and environmental information. Philos. Trans. R. Soc. B Biol. Sci., 370, 20140016.

Jetz, W., Thomas, G.H., Hartmann, K. & Mooers, A.O. (2012). The global diversity of birds in space and time. Nature, 491, 444–448.

Kembel, S.W., Cowan, P.D., Helmus, M.R., Cornwell, W.K., Morlon, H., Ackerly, D.D., et al. (2010). Picante: R tools for integrating phylogenies and ecology. Bioinformatics, 26, 1463–1464.

Kline, R.B. (2010). Principles and practice of structural equation modeling. 3rd edn. The Guilford Press, New York, NY, USA.

De Laender, F., Rohr, J.R., Ashauer, R., Baird, D.J., Berger, U., Eisenhauer, N., et al. (2016). Reintroducing environmental change drivers in biodiversity–ecosystem functioning research. Trends Ecol. Evol., 31, 905–915.

Laliberté, E. & Legendre, P. (2010). A distance-based framework for measuring functional diversity from multiple traits. Ecology, 91, 299–305.

Laliberté, E., Legendre, P. & Shipley, B. (2015). FD: measuring functional diversity from multiple traits, and other tools for functional ecology. R package version 1.0-12. Available for download from: https://cran.r-project.org/package=FD.

Lamb, E.G., Mengersen, K.L., Stewart, K.J., Attanayake, U. & Siciliano, S.D. (2014). Spatially explicit structural equation modeling. Ecology, 95, 2434–2442.

Lee, T.M. & Jetz, W. (2011). Unravelling the structure of species extinction risk for predictive conservation science. Proc. R. Soc. B Biol. Sci., 278, 1329–1338.

Levine, J.M., Adler, P.B. & Yelenik, S.G. (2004). A meta-analysis of biotic resistance to exotic plant invasions. Ecol. Lett., 7, 975–989.

Mittelbach, G.G., Schemske, D.W., Cornell, H. V., Allen, A.P., Brown, J.M., Bush, M.B., et al. (2007). Evolution and the latitudinal diversity gradient: Speciation, extinction and biogeography. Ecol. Lett., 10, 315–331.

Naeem, S., Knops, J.M., Tilman, D., Howe, K.M., Kennedy, T. & Gale, S. (2000). Plant diversity increases resistance to invasion in the absence of covarying extrinsic factors. Oikos, 91, 97–108.

Naeem, S., Prager, C., Weeks, B.C., Varga, A., Flynn, D., Griffin, K., et al. (2016). Biodiversity as a multidimensional construct. Proc. R. Soc. B Biol. Sci., 283, 20153005.

Oliver, T.H., Heard, M.S., Isaac, N.J.B., Roy, D.B., Procter, D., Eigenbrod, F., et al. (2015). Biodiversity and resilience of ecosystem functions. Trends Ecol. Evol., 30, 673–684.

Owens, I.P.F., Bennett, P.M. & Harvey, P.H. (1999). Species richness among birds: Body size, life history, sexual selection or ecology? Proc. R. Soc. B Biol. Sci., 266, 933–939.

Pigot, A.L., Sheard, C., Miller, E.T., Bregman, T.P., Freeman, B.G., Roll, U., et al. (2020). Macroevolutionary convergence connects morphological form to ecological function in birds. Nat. Ecol. Evol., 4, 230–239.

Pigot, A.L., Tobias, J.A. & Jetz, W. (2016). Energetic constraints on species coexistence in birds. PLoS Biol., 14, e1002407.

R Core Team. (2018). R: A language and environment for statistical computing. Available for download from: https://cran.r-project.org.

Rabinowitz, D. (1981). Seven forms of rarity. In: The biological aspects of rare plant conservation. Wiley, Chichester, UK, pp. 205–217.

Reich, P.B., Tilman, D., Isbell, F., Mueller, K., Hobbie, S.E., Flynn, D.F.B., et al. (2012). Impacts of biodiversity loss escalate through time as redundancy fades. Science, 336, 589–592.

Reinhardt, K., Kohler, G., Maas, S. & Detzel, P. (2005). Low dispersal ability and habitat specificity promote extinctions in rare but not in widespread species: the Orthoptera of Germany. Ecography, 28, 593–602.

Rosseel, Y. (2012). lavaan : an R package for structural equation modeling. J. Stat. Softw., 48, 1–36.

Rubolini, D., Liker, A., Garamszegi, L.Z., Møller, A.P. & Saino, N. (2015). Using the birdtree.org website to obtain robust phylogenies for avian comparative studies: a primer. Curr. Zool., 61, 959–965.

Şekercioǧlu, Ç.H., Daily, G.C. & Ehrlich, P.R. (2004). Ecosystem consequences of bird declines. Proc. Natl. Acad. Sci. U. S. A., 101, 18042–18047.

Sheard, C., Neate-Clegg, M.H.C., Alioravainen, N., Jones, S.E.I., Vincent, C., MacGregor, H.E.A., et al. (2020). Ecological drivers of global gradients in avian dispersal inferred from wing morphology. Nat. Commun., 11.

Soliveres, S., Van Der Plas, F., Manning, P., Prati, D., Gossner, M.M., Renner, S.C., et al. (2016). Biodiversity at multiple trophic levels is needed for ecosystem multifunctionality. Nature, 536, 456–459.

Sukumaran, J. & Holder, M.T. (2010). DendroPy: A Python library for phylogenetic computing. Bioinformatics, 26, 1569–1571.

Tilman, D., Isbell, F. & Cowles, J.M. (2014). Biodiversity and ecosystem functioning. Annu. Rev. Ecol. Evol. Syst., 45, 471–493.

Tilman, D., Knops, J., Wedin, D., Reich, P., Ritchie, M. & Siemann, E. (1997). The influence of functional diversity and composition on ecosystem processes. Science, 277, 1300–1302.

Tobias, J.A. AVONET: A global database of bird traits. Ecol. Lett. Submitted for publication in the same special section within this issue.

Tobias, J.A., Ottenburghs, J. & Pigot, A.L. (2020). Avian diversity: speciation, macroevolution and ecological function. Annu. Rev. Ecol. Evol. Syst., 51.

Tobias, J.A. & Pigot, A.L. (2019). Integrating behaviour and ecology into global biodiversity conservation strategies. Philos. Trans. R. Soc. B Biol. Sci., 374, 20190012.

Villéger, S., Mason, N.W.H. & Mouillot, D. (2008). New multidimensional functional diversity indices for a multifaceted framework in functional ecology. Ecology, 89, 2290–2301.

Webb, C.O., Ackerly, D.D., McPeek, M.A. & Donoghue, M.J. (2002). Phylogenies and community ecology. Ann. Rev. Ecol. Evol. Syst., 33, 475–505.

Weeks, B.C. & Claramunt, S. (2014). Dispersal has inhibited avian diversification in Australasian archipelagoes. Proc. R. Soc. B Biol. Sci., 281, 20141257.

Weeks, B.C., Claramunt, S. & Cracraft, J. (2016a). Integrating systematics and biogeography to disentangle the roles of history and ecology in biotic assembly. J. Biogeogr., 43, 1546–1559.

Weeks, B.C., Gregory, N. & Naeem, S. (2016b). Bird assemblage vulnerability depends on the diversity and biogeographic histories of islands. Proc. Natl. Acad. Sci. U. S. A., 113, 10109–10114.

Wilman, H., Belmaker, J., Simpson, J., de la Rosa, C., Rivadeneira, M.M. & Jetz, W. (2014). EltonTraits 1.0: Species-level foraging attributes of the world’s birds and mammals. Ecology, 95, 2027–2027.

Winemiller, K.O., Fitzgerald, D.B., Bower, L.M. & Pianka, E.R. (2015). Functional traits, convergent evolution, and periodic tables of niches. Ecol. Lett., 18, 737–751.

